# Population divergence in growth allometry of annual killifishes across ephemeral habitats in Malawi floodplain systems

**DOI:** 10.64898/2026.07.21.739734

**Authors:** Felix Sanudi, Fanuel Kapute, Benjamin Kondowe, Kumbukani Mzengereza, Wilson Sawasawa, Geoffrey Kanyerere, Maggie Munthali, Emmanuel Cishibanji, Wales Singini, Enoch Ng’oma

## Abstract

Body size is a fundamental determinant of ecological performance, and variation in length-weight relationships can provide insight into how populations respond to local environmental conditions. Annual killifishes inhabit highly seasonal wetlands characterized by substantial environmental heterogeneity, yet population-level variation in growth allometry remains poorly understood. We investigated growth allometry in 18 populations of two annual killifish species, *Nothobranchius kirki* and *N. wattersi*, distributed across ephemeral habitats in Malawi. Length-weight relationships were analyzed separately for each species using linear mixed-effects models, with locality incorporated as a random effect. Environmental variation among localities was summarized using principal component analysis of water quality variables, and population-specific allometric coefficients were subsequently related to environmental gradients. Significant population-level variation in growth trajectories was detected in both species, indicating divergence in allometric scaling among localities. Divergence was more pronounced in *N. wattersi*, which exhibited a broader range of population-specific allometric coefficients than *N. kirki*. Populations also differed significantly in relative body condition after accounting for body length and sex. Environmental gradients explained a significant proportion of variation in allometric slopes in *N. kirki*, whereas no significant relationship was detected in *N. wattersi*. Thus, the species exhibiting weaker allometric divergence showed stronger environmental associations, while the species exhibiting greater divergence showed little correspondence with measured environmental variables. These results demonstrate substantial spatial heterogeneity in growth allometry among populations of annual killifishes inhabiting seasonal wetlands. Furthermore, the contrasting environmental associations observed between species suggest that population divergence in growth trajectories may arise through different ecological and evolutionary processes, even among closely related taxa occupying similar habitats.

## Introduction

Body size is a fundamental biological trait that influences virtually every aspect of organismal ecology, including metabolism, resource acquisition, reproductive output, and survival (Elton, 1927; Kleiber, 1932; Senthilnathan, 2023). In fishes, the relationship between body length and body mass provides a widely used framework for quantifying growth patterns and evaluating variation in body condition among individuals and populations (Audzijonyte *et al*., 2025; Kamble *et al*., 2024). Length-weight relationships (LWR) have therefore become an important tool for understanding how environmental and ecological factors influence growth dynamics across spatial and temporal scales (Audzijonyte *et al*., 2025; Froese, 2006; Le Cren, 1951). Although many fish populations exhibit approximately isometric growth (Kumari *et al*., 2025; Ragheb, 2023), deviations from isometry are common (Bolger & Connolly, 1989; Froese, 2006) and may reflect differences in resource availability, habitat quality, population density, environmental stress, or life-history strategy (Miranda *et al*., 2026).

Spatial variation in allometric growth has attracted increasing attention because populations occupying different environments often experience distinct ecological conditions that can influence energy acquisition and allocation (Boukal *et al*., 2014; Dmitriew, 2011; Ylikarjula *et al*., 1999). Environmental factors such as temperature, dissolved oxygen, salinity, conductivity, hydroperiod, and productivity have been shown to affect growth rates and body condition in fishes (Duan, 1998; Le Cren, 1951; López-Pérez *et al*., 2020). Consequently, variation in length-weight scaling among populations may provide insight into how local environments shape growth trajectories. However, environmental influences on growth are frequently complex, and population differences may arise not only from contemporary environmental conditions but also from phenotypic plasticity, local adaptation, or interactions among multiple ecological factors (Dikou, 2023; Dmitriew, 2011).

Annual killifishes (genus *Nothobranchius* Peters) provide an excellent system for investigating population-level variation in growth allometry. These fishes inhabit temporary wetlands and ephemeral pools that form during seasonal rains and persist for only a few months before drying completely (Furness, 2016; Terzibasi Tozzini & Cellerino, 2020). Because their aquatic habitat is highly time constrained, annual killifishes have evolved rapid growth, early maturation, and accelerated reproduction, enabling completion of their life cycle within a single wet season (Cellerino *et al*., 2016; Furness, 2016). Their dependence on short-lived habitats exposes populations to substantial spatial heterogeneity in environmental conditions, including differences in water chemistry, hydroperiod, vegetation structure, and resource availability (Furness, 2016; Polačik & Reichard, 2010). Such environmental variation has the potential to generate differences in growth trajectories among populations despite their shared life-history strategy.

In this study, we focus on two allopatric species of annual killifish in Malawi, *N. kirki* and *N. wattersi*, distributed across distinct floodplain systems associated with Lake Malawi and the Lake Chilwa-Chiuta basin (Ng’oma *et al*., 2013). These floodplains comprise networks of seasonal wetlands that differ in hydrological and physicochemical characteristics, potentially creating contrasting growth environments for local populations. Despite growing interest in the ecology and evolution of annual killifishes (Cellerino *et al*., 2016; Dorn *et al*., 2011; Terzibasi Tozzini & Cellerino, 2020), quantitative evaluations of growth allometry across multiple wild populations remain limited. As a result, the extent to which population-level variation in growth trajectories reflects local environmental conditions remains poorly understood.

Here, we examined length-weight relationships in 18 natural populations of *N. wattersi* and *N. kirki* distributed across Malawi floodplain habitats. Specifically, we asked whether growth allometry varies among populations and whether population differences are associated with environmental gradients. We predicted that (i) growth trajectories would differ among populations, reflecting spatial variation in ecological conditions across ephemeral habitats, and (ii) environmental variables would explain a portion of the observed variation in allometric scaling among populations. By integrating mixed-effects modelling with environmental analyses, this study provides one of the first assessments of population-level variation in growth allometry in wild African annual killifishes and contributes to a broader understanding of how environmental heterogeneity influences growth patterns in seasonal aquatic systems.

## Materials and Methods

### Study system and sampling

A total of 824 annual killifishes representing two species, *N. wattersi* and *N. kirki*, were collected from ephemeral pools within two major floodplain systems in Malawi during the 2024 and 2025 rainy seasons. Sampling included 620 individuals of *N. wattersi* (295 males and 325 females) from nine populations distributed across the Lake Malawi floodplain and 204 individuals of *N. kirki* (101 males and 103 females) from nine populations within the Lake Chilwa-Chiuta floodplain (Fig.1). Fish were collected using hand-held dip nets and small seine nets (4-5 m length) in temporary pools, seasonally inundated wetlands and rice paddies. Each specimen was identified to species, sexed, and measured for total length (TL, mm) using digital calipers (General^®^ UltraTech^®)^. Body mass (g) was measured using an electronic pocket scale (Nutri Fit, model: WEHA0151H). Environmental conditions at each locality were characterized using in situ measurements of water temperature (°C), dissolved oxygen (mg L⁻¹), conductivity (mS cm⁻¹), salinity (PSU), and pH using portable probes (Multiparameter model HI98194, Hanna Instruments).

**Figure 1.**
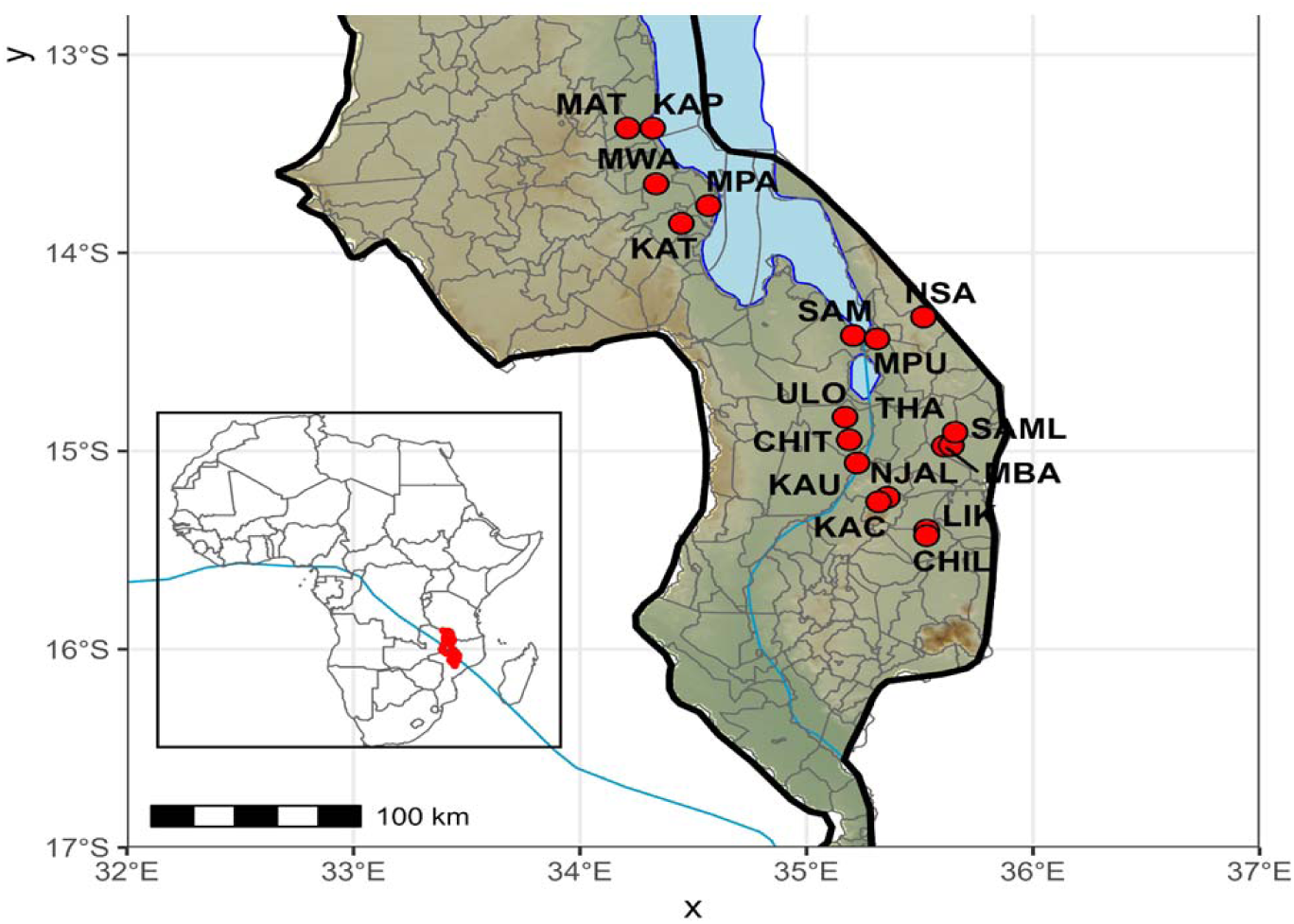
Map of Malawi with an inset of Africa showing sampling locations. Points indicate sampling location in the Lake Malawi and Lake Chilwa/Chiuta floodplains.

### Length-weight relationships and growth allometry

Population variation in growth allometry was evaluated using the length-weight relationship. Body mass and total length were log-transformed prior to analysis to linearize the allometric relationship and stabilize variance. Analyses were conducted separately for *N. wattersi* and *N. kirki* to account for species-specific differences in morphology and ecology. For each species, linear mixed-effects models (LMMs) were fitted using the package lme4 in R (R Core Team). Log-transformed body mass was modeled as a function of centered log-transformed body length and sex:

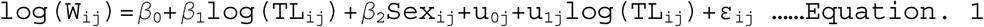

where β_0_ is the fixed intercept, β_1_ is the average allometric coefficient relating body length to body mass, β_2_ represents the fixed effect of sex, u_0j_ is the locality-specific random intercept, u_1j_is the locality-specific random slope for body length, and ε_ij_ is the residual error term. Subscripts *`i`* and *j* denote individual fish and locality, respectively.

To evaluate spatial variation in growth trajectories, locality was incorporated as a random effect. To test whether growth trajectories differed among populations, we compared a random-intercept model with a random-slope model using likelihood-ratio tests. The random-intercept model included locality as a random intercept only, whereas the random-slope model additionally allowed the relationship between body length and body mass to vary among localities. Centering of log-transformed length was performed prior to analysis to improve model convergence and facilitate interpretation of model parameters. The fixed effect of body length represents the average allometric scaling relationship across populations, whereas locality-specific random slopes quantify deviations from the overall growth trajectory among populations.

### Population divergence in growth trajectories

To test whether growth trajectories differed among populations, two competing mixed-effects models were compared for each species. The first model included locality as a random intercept only, whereas the second included both random intercepts and locality-specific random slopes. Models were compared using likelihood ratio tests (LRTs). A significant improvement in model fit following inclusion of random slopes was interpreted as evidence that the relationship between body length and body mass differed among populations, indicating population-level divergence in growth allometry. Model performance was assessed using marginal and conditional coefficients of determination (R^2^) following Nakagawa and Schielzeth (2013). Marginal R^2^ describes variation explained by fixed effects alone, whereas conditional R^2^ represents variation explained by the combined effects of fixed and random components.

Population-specific allometric coefficients were extracted from the fitted mixed models and used to visualize variation in growth trajectories among localities.

### Relative body condition

To evaluate variation in relative body condition among populations, an analysis of covariance (ANCOVA) was performed using log-transformed body mass as the response variable, log-transformed body length as a covariate, sex as a fixed effect, and locality as a factor. Estimated marginal means were calculated at the mean body length to compare length-adjusted body mass among populations. Pairwise differences among localities were assessed using Tukey-adjusted comparisons.

### Environmental variation among localities

To characterize environmental variation among sampling sites, principal component analyses (PCA) were conducted separately for *N. wattersi* and *N. kirki* using standardized environmental variables. Analyses included temperature, dissolved oxygen, conductivity, salinity, and pH. These environmental variables were centered and scaled prior to PCA. Principal components explaining the largest proportion of environmental variation were retained for interpretation. Variable loadings were examined to identify environmental gradients represented by individual principal component axes.

### Environmental correlates of allometric variation

To evaluate whether environmental variation contributed to population differences in growth allometry, locality-specific allometric slopes extracted from the mixed-effects models were regressed against environmental principal component scores. Separate linear models were fitted for each species using population-specific allometric slopes as the response variable and the first two principal component axes as predictors:

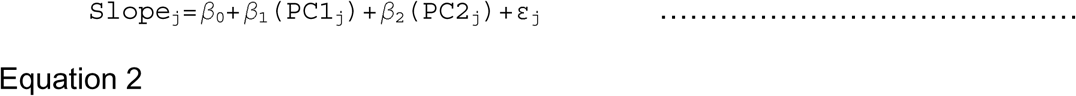

where Slope_j_ represents the locality-specific allometric coefficient estimated from the mixed-effects model, PC1 and PC2 are the first two environmental principal component axes, and ε_j_ is the residual error term. Model significance was evaluated using F-tests, and the contribution of individual environmental axes was assessed using regression coefficients and associated probability values. Significant relationships were interpreted as evidence that environmental gradients were associated with variation in population-level growth trajectories.

### Statistical analyses

All analyses were performed in R version 4.6.0 (R Core Team). Linear mixed-effects models were fitted using the package lme4, model comparisons were conducted using likelihood ratio tests, coefficients of determination were calculated using performance, and principal component analyses were performed using `baseR` functions. Statistical significance was assessed at α = 0.05.

## Results

### Length-weight relationships

Length and body mass exhibited strong positive allometric relationships in both species. Body mass increased predictably with body length in both species, although the relationship varied among localities (Fig. 2). In mixed effects models, log-transformed body length was a highly significant predictor of log-transformed body mass in both *N. kirki* (β = 3.123 ± 0.104, t = 30.09) and *N. wattersi* (β = 3.039 ± 0.158, t = 19.19). Sex did not significantly influence body mass in *N. kirki* (β = - 0.013 ± 0.019, t = - 0.68), whereas a small positive effect of male sex was detected in *N. wattersi* (β = 0.032 ± 0.011, t = 3.03). Mixed effects models explained a large proportion of variation in body mass for both species. Marginal and conditional coefficients of determination were high in *N. kirki* (R^2^_m_ = 0.961, R^2^ = 0.976) and *N. wattersi* (R^2^ = 0.929, R^2^ = 0.968), indicating that body length accounted for most variation in body mass while locality effects contributed additional explanatory power.

**Figure 2.**
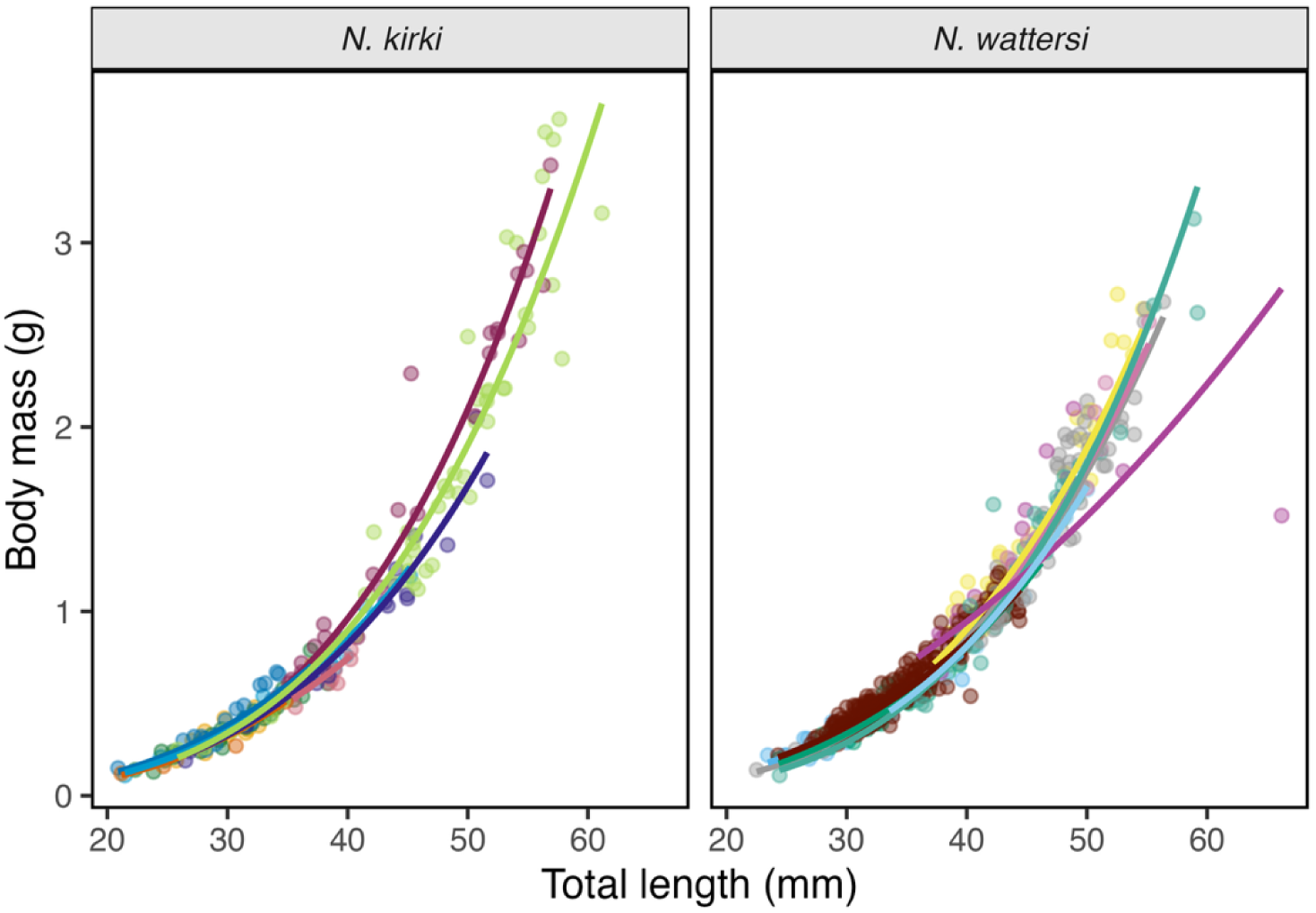
Length-weight relationships for populations of *N. kirki* and *N. wattersi* sampled from seasonal pools in Malawi. Points represent individual fish and curves represent locality-specific predictions from mixed effects allometric models. Predictions were generated from models fitted to log-transformed body mass and body length and subsequently back-transformed to the original scale. Colors indicate sampling localities.

### Population divergence in growth trajectories

Growth trajectories differed significantly among populations in both species. Likelihood-ratio tests comparing random intercept and random slope models demonstrated that allowing locality-specific allometric slopes significantly improved model fit (Table 1). For *N. kirki*, the inclusion of locality-specific slopes produced a significant improvement in model fit (χ² = 8.85, df = 2, p = 0.012), indicating that populations differed in the relationship between body length and body mass. Similar but substantially stronger evidence was observed in *N. wattersi*, where random slope models outperformed random intercept models (χ² = 71.43, df = 2, p < 0.001).

**Table 1.** Summary of models examining population divergence in growth allometry and relative body condition in *N. kirki* and *N. wattersi.* Columns report fixed-effect coefficients from mixed-effects length-weight models, likelihood-ratio tests comparing random intercept and random slope models, marginal (R²_m_) and conditional (R²_c_) coefficients of determination, tests of population differences in relative body condition, and regressions of locality-specific allometric slopes against environmental principal component axes

| Species | Growth coefficient<br>($\beta \pm \text{SE}$) | Random slope<br>test $\chi^2$ (p) | $R^2_m$ \| $R^2_c$ | Condition<br>F (p) | Environment<br>allometry | |
| --- | --- | --- | --- | --- | --- | --- |
| | | | | | $R^2$ | F (p) |
| *N. kirki* | 3.123 ± 0.104 | 8.85 (0.012) | 0.961 \| 0.976 | 8.33<br>($<0.001$) | 0.726 | 7.93 (0.021) |
| *N. wattersi* | 3.039 ± 0.158 | 71.43 ($<0.001$) | 0.929 \| 0.968 | 11.23<br>($<0.001$) | 0.182 | 0.67 (0.548) |

Population-specific allometric coefficients varied substantially among localities, ranging from 2.9 to 3.4 in *N. kirki* and from 2.2 to 3.6 in *N. wattersi* (Fig. 3). The variance associated with locality-specific slopes was greater in *N. wattersi* (σ² = 0.202) than in *N. kirki* (σ² = 0.062), suggesting stronger population level divergence in growth trajectories within the Lake Malawi floodplain system. Together, these results indicate that growth allometry is not uniform across populations but instead varies spatially within both species.

**Figure 3.**
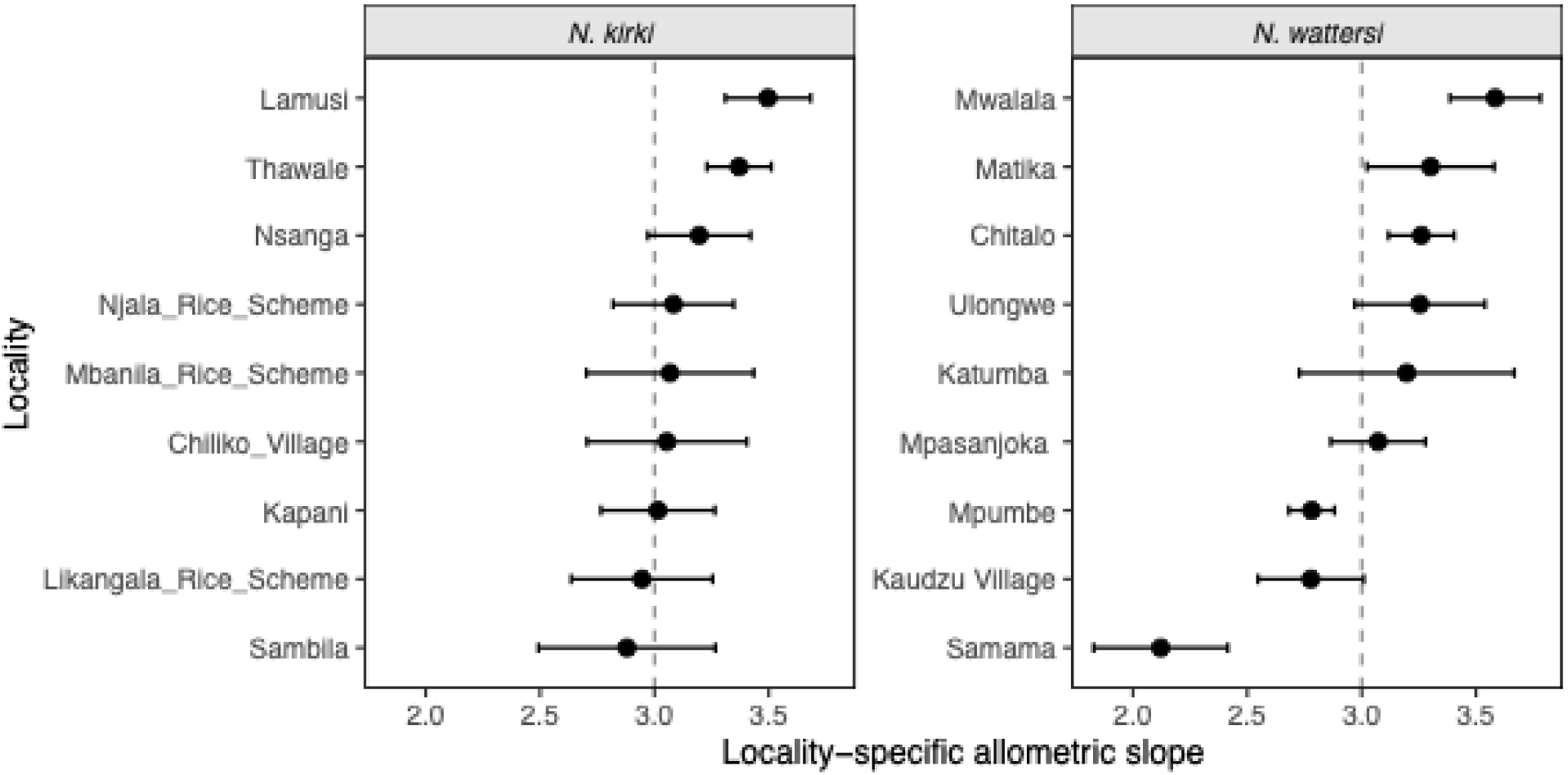
Allometric coefficients for populations estimated from mixed effects length-weight models for *N. kirki* and *N. wattersi*. Points represent allometric slope for each locality (β₁) describing the relationship between log-transformed body mass and log-transformed body length, and horizontal bars indicate 95% confidence intervals. The dashed line denotes the species-wide mean allometric coefficient estimated from the model. Localities are ordered by increasing slope within each species. Variation among localities indicates divergence in growth trajectories at the population level.

### Population variation in relative body condition

Relative body condition differed significantly among populations in both species after accounting for body length and sex (Table 1, Table S1). In *N. wattersi*, locality exerted a strong effect on length-adjusted body mass (F_8,1032_ = 11.23, p < 0.001). Estimated marginal means indicated that individuals from Matika and Katumba exhibited the highest relative condition, whereas those from Mpasanjoka and Ulongwe had the lowest (Fig. S1). Similar patterns were observed in *N. kirki* (F_8,193_ = 8.33, p < 0.001), with the highest length-adjusted body mass recorded in Lamusi and Likangala Rice Scheme and the lowest in Sambila, Mbanila Rice Scheme, and Nsanga. Residual condition analyses yielded qualitatively identical results, confirming substantial spatial variation in body condition within both floodplain systems. Further, population-specific allometric slopes showed weak associations with relative body condition in both species (Fig. S2).

### Environmental variation among localities

Environmental conditions varied among sampling localities in both species (Fig. 4). Principal component analysis of environmental variables revealed that the first two axes accounted for 65.8% of total environmental variation in *N. kirki* (PC1 = 44.7%, PC2 = 21.1%) and 69.3% of total variation in *N. wattersi* (PC1 = 43.0%, PC2 = 26.3%). In *N. kirki*, PC1 represented a broad physicochemical gradient, with the strongest loadings associated with pH and temperature, followed by salinity and conductivity. PC2 was influenced primarily by salinity and dissolved oxygen. In *N. wattersi*, PC1 was driven mainly by temperature and dissolved oxygen, with additional contributions from pH and conductivity, whereas PC2 was associated primarily with salinity and conductivity. Together, these axes captured the major environmental gradients distinguishing floodplain habitats within each species. With respect to body condition, environmental principal component axes showed limited relationships with relative body condition (Fig. S3).

**Figure 4.**
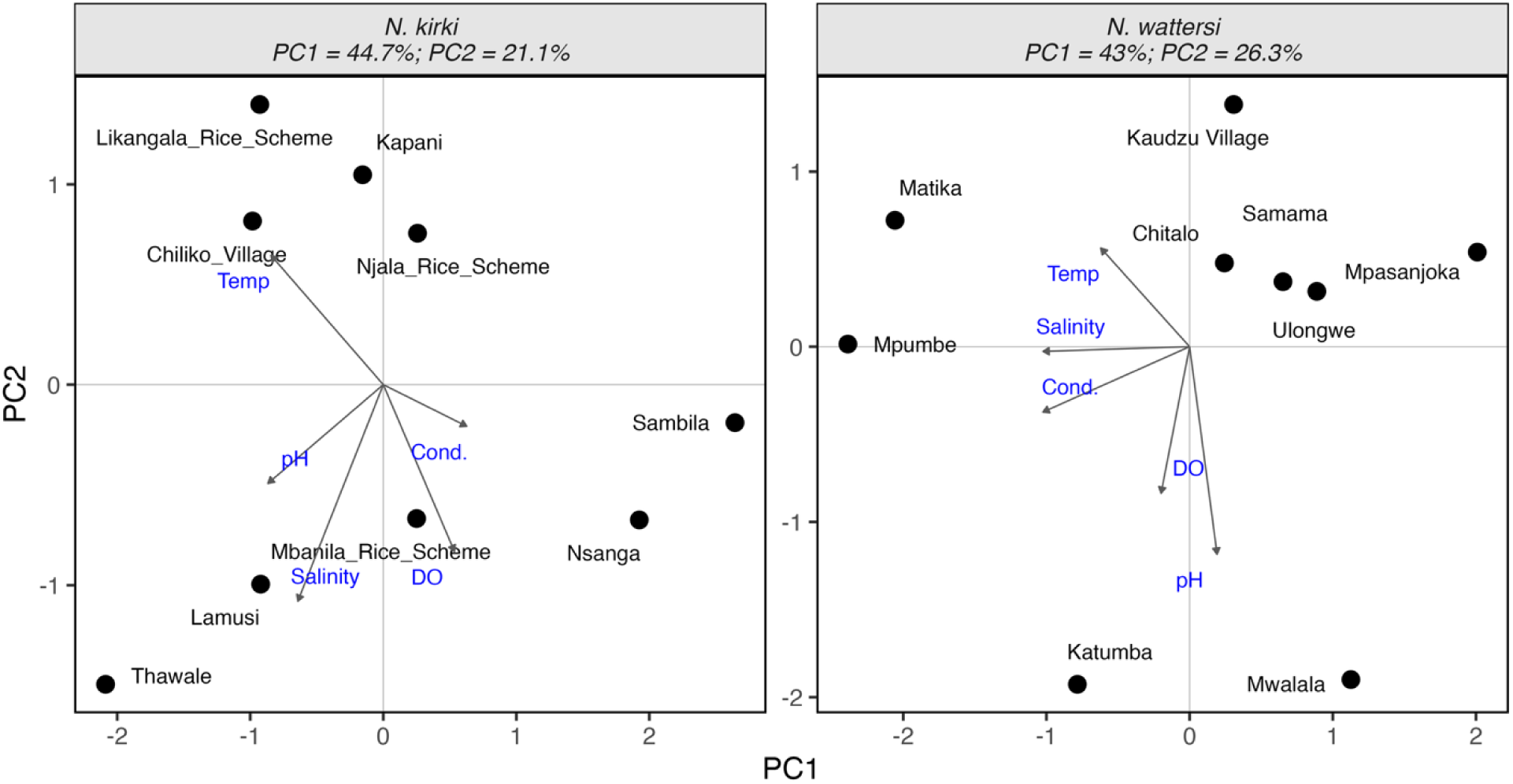
Principal component analysis of environmental variation among sampling localities for *N. kirki* and *N. wattersi*. Points represent locality scores based on standardized environmental variables, and arrows indicate variable loadings for temperature, salinity, conductivity, pH, and dissolved oxygen. PCA axes summarize environmental gradients used to test associations between local habitat conditions and population allometric slopes.

### Environmental correlates of allometric variation

Environmental variables explained significant variation in locality-specific allometric slopes in *N. kirki* but not in *N. wattersi* (Fig. 5; Table 1). In *N. kirki*, allometric slopes were significantly associated with environmental variation (F_2,6_ = 7.93, p = 0.021, R^2^ = 0.726). PC2 exhibited a significant negative relationship with allometric slope (β = - 0.137 ± 0.041, p = 0.017), while PC1 showed a marginally significant negative association (β = - 0.064 ± 0.028, p = 0.067). Together, the first two environmental axes explained approximately 73% of variation in population-level growth trajectories. In contrast, environmental variables explained little variation in growth allometry among populations of *N. wattersi* (F_2,6_ = 0.67, p = 0.548, R^2^ = 0.182). Neither PC1 (β = 0.011 ± 0.108, p = 0.919) nor PC2 (β = - 0.158 ± 0.138, p = 0.294) was significantly associated with locality-specific allometric slopes. These results indicate that environmental gradients were associated with population differences in growth allometry in *N. kirki*, whereas comparable relationships were not detected in *N. wattersi*.

**Figure 5.**
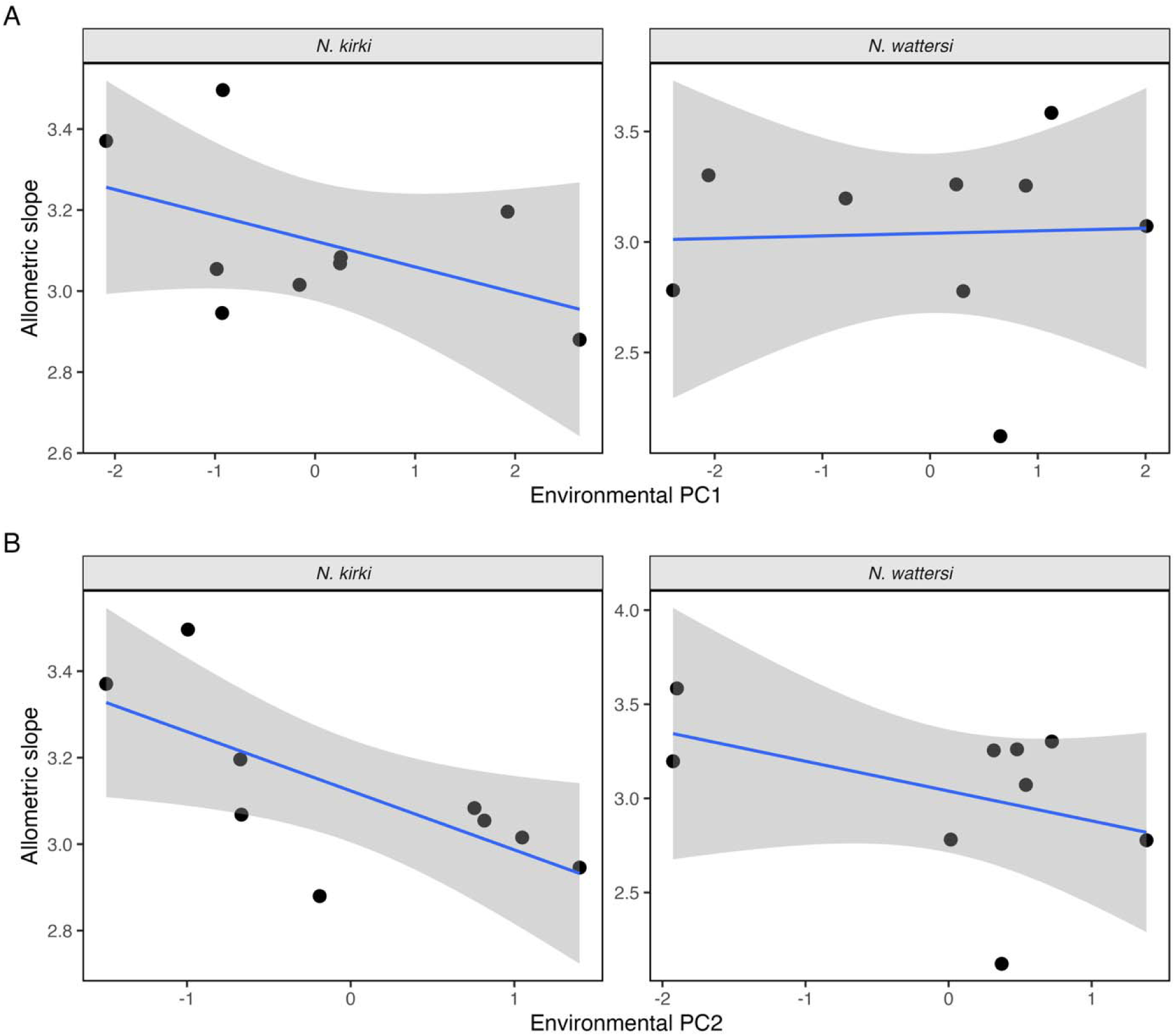
Relationships between environmental gradients and allometric coefficients in populations of *N. kirki* and *N. wattersi*. Allometric slopes were estimated from mixed effects length-weight models and regressed against environmental principal component axes derived from standardized measurements of temperature, salinity, conductivity, dissolved oxygen, and pH. **(a)** Relationships between allometric slope and environmental PC1, **(b)** relationships with environmental PC2. Points represent localities, blue lines indicate fitted linear regressions with 95% confidence intervals in grey shade.

## Discussion

Body size is a fundamental determinant of ecological performance in fishes, influencing metabolism, resource acquisition, reproductive output, and survival (Froese, 2006; Kleiber, 1932). Consequently, variation in length-weight relationships can provide insight into how populations respond to local environmental conditions and how growth trajectories diverge across landscapes. In this study, we examined growth allometry in wild populations of two annual killifishes, *Nothobranchius kirki* and *N. wattersi*, distributed across seasonal habitats in Malawi. Our analyses revealed significant population-level variation in growth trajectories within both species, demonstrating that growth allometry is not uniform across the landscape despite the shared annual life-history strategy characteristic of the genus. Population differences were accompanied by variation in relative body condition, and environmental gradients were associated with allometric divergence in *N. kirki* but not in *N. wattersi*. Our results indicate that growth patterns in annual killifish exhibit substantial spatial heterogeneity and that the mechanisms generating this variation differ between species.

### Population divergence in growth allometry

The key result emerging from this study was the presence of significant population-level variation in allometric scaling within both species. Although body mass increased predictably with body length in all populations, the rate at which mass accumulated with increasing length differed among localities. Such variation indicates that populations do not simply differ in average body size but instead follow distinct growth trajectories. Differences in allometric scaling are ecologically important because they influence energetic demands, reproductive allocation, swimming performance, and overall body condition, thereby affecting multiple components of fitness (Froese, 2006; Le Cren, 1951). Spatial variation in length-weight relationships has been documented across a wide range of fish taxa and is frequently attributed to differences in habitat quality, resource availability, environmental conditions, and population density (Blackwell *et al*., 2000; Bolger & Connolly, 1989; Froese, 2006). However, most studies focus on variation in average condition or population-specific length-weight parameters rather than explicitly testing whether populations differ in their allometric trajectories. Our results therefore extend previous work by demonstrating that populations of annual killifish occupying distinct floodplain systems exhibit significant divergence in the relationship between body length and body mass.

The magnitude of divergence differed substantially between species. Although significant variation in allometric slopes was detected in both *N. kirki* and *N. wattersi*, divergence was considerably stronger in *N. wattersi*, where locality-specific slopes spanned a broader range and inclusion of random slopes greatly improved model fit. This pattern suggests that populations of *N. wattersi* experience greater ecological or evolutionary heterogeneity than populations of *N. kirki*. Similar patterns of population-level divergence in growth-related traits have been reported in other fishes and are often interpreted as evidence of local adaptation, environmentally induced plasticity, or interactions between the two processes (Conover *et al*., 2009; Fraser *et al*., 2011; Taylor, 1991). While our analyses do not allow discrimination among these alternative interpretations, they demonstrate that substantial growth divergence can arise even among populations sharing a broadly similar annual life-history strategy.

### Variation in relative body condition among populations

In addition to differences in growth trajectories, populations differed significantly in relative body condition after accounting for body length and sex. These differences indicate that fish from some localities were consistently heavier or lighter than expected for a given length, suggesting variation in energy reserves, nutritional status, growth efficiency, or recent environmental history among populations (Bolger & Connolly, 1989; Le Cren, 1951). Variation in body condition is widely understood as an indicator of habitat quality and population performance because it integrates multiple ecological influences, including food availability, physiological stress, competition, and reproductive investment (Blackwell *et al*., 2000). Importantly, divergence in relative body condition and divergence in allometric scaling represent distinct aspects of population variation. Relative condition reflects differences in body mass after accounting for length, whereas allometric scaling describes how body mass changes with increasing body length across the size range of a population (Bolger & Connolly, 1989; Froese, 2006; Le Cren, 1951). Consequently, populations may differ in average mass at a given length while exhibiting similar growth trajectories, or alternatively may display divergent allometric relationships despite comparable body condition. The coexistence of both patterns in our analyses suggests that spatial variation in growth among annual killifish populations occurs through multiple pathways, influencing both the shape of the length-weight relationship and the average body mass attained by individuals.

### Environmental correlates of growth divergence

A central objective of this study was to determine whether environmental variation among floodplains contributes to population-level divergence in growth allometry. Environmental conditions influence fish growth through multiple physiological pathways, including effects on metabolic rate, oxygen availability, osmoregulation, and energy allocation (Brett; Claireaux & Chabot, 2016; Pörtner & Knust, 2007). Because annual killifishes inhabit environmentally variable temporary pools, we predicted that local environmental conditions would be associated with variation in growth trajectories. Consistent with this prediction, environmental principal components explained a significant proportion of variation in allometric slopes across localities in *N. kirki*. In particular, the second environmental axis was significantly associated with variation in growth trajectories, indicating that populations occupying environmentally distinct habitats tended to exhibit different patterns of length-weight scaling. Although PCA axes integrate multiple environmental variables and therefore cannot be interpreted as single factor effects, this result nevertheless suggests that contemporary environmental conditions contribute to shaping growth dynamics in *N. kirki*. Similar relationships between environmental gradients and growth-related traits have been reported across diverse fish systems, where variation in temperature, oxygen availability, salinity, and habitat productivity influences growth performance and energy allocation (Brett; Pörtner & Knust, 2007).

In contrast, environmental gradients did not explain variation in allometric slopes in *N. wattersi*. This result is particularly noteworthy because *N. wattersi* exhibited stronger overall divergence in growth allometry than *N. kirki*. Thus, the species showing the greatest population differentiation was also the species in which measured environmental variation provided the least explanatory power. This contrast suggests that different processes may underlie growth divergence in the two species. One possibility is that the environmental variables measured here capture the principal ecological drivers affecting growth in *N. kirki* but not in *N. wattersi*. Factors such as hydroperiod duration, prey abundance, vegetation structure, predation pressure, or population density may differ among *N. wattersi* localities yet remain uncaptured by the environmental dataset used in this study. Alternatively, growth divergence in *N. wattersi* may reflect stronger contributions from historical or evolutionary processes. Population differentiation often arises through a combination of environmental variation, phenotypic plasticity, and local adaptation (Fraser *et al*., 2011; Ghalambor *et al*., 2007; West-Eberhard, 2003). If *N. wattersi* populations have experienced longer-term isolation or stronger divergent selection, differences in growth trajectories may persist even when contemporary environmental variables show limited explanatory power.

The contrasting results between species therefore represent one of the most informative outcomes of this study. Rather than suggesting a single mechanism underlying growth divergence in annual killifish, our findings indicate that the relative importance of environmental and potentially evolutionary influences may vary among closely related taxa occupying similar habitats. Future studies integrating genomic, common garden, or reciprocal transplant approaches could help distinguish environmentally induced plasticity from genetically based population divergence.

### Implications for annual killifish ecology

Annual killifishes are often viewed as ecological specialists constrained by the extreme seasonality of temporary pools. Because these habitats persist for only a few months before drying completely, strong selection favors rapid growth, early maturation, and accelerated reproduction (Cellerino *et al*., 2016; Genade *et al*., 2005). Under such constraints, one might expect growth trajectories to be relatively conserved among populations. Instead, our results demonstrate substantial variation within species in both growth allometry and body condition.

These findings indicate that the annual life history strategy does not impose a single invariant growth pattern. Rather, populations appear capable of achieving the same fundamental life-history pattern (i.e. completion of the life cycle within a limited hydroperiod) through different growth trajectories. Such flexibility is consistent with broader evidence that annual killifishes occupy environmentally heterogeneous habitats and exhibit considerable ecological variation across their geographic ranges (Cellerino *et al*., 2016; Reichard *et al*., 2009). Our results therefore reinforce the view that annual killifish represent highly dynamic systems in which local environmental conditions and population history can interact to generate substantial biological diversity.

## Conclusions

Our study demonstrates that populations of *Nothobranchius kirki* and *N. wattersi* exhibit significant divergence in growth allometry across ephemeral floodplain habitats in Malawi. Population differences were evident not only in growth trajectories but also in relative body condition, indicating substantial spatial heterogeneity in growth dynamics within both species. Most importantly, environmental gradients explained variation in allometric slopes in *N. kirki* but not in *N. wattersi*, suggesting that the drivers of growth divergence differ between species. These findings reveal that growth allometry in annual killifishes is more variable than previously recognized and highlight the importance of considering both environmental and evolutionary processes when interpreting spatial variation in growth patterns across seasonal aquatic ecosystems.

## Acknowledgements

We would like to acknowledge the communities around Lake Malawi and the Lake Chilwa/Chiuta floodplains for providing guidance and helping identify pools for our sampling. We also thank the Department of Fisheries and Aquatic Sciences at Mzuzu University for hosting the Mzuzu University – University of Missouri Killifish Project, which made this study possible.

## Author contributions

F.S. generated the study idea, collected field samples, performed statistical analysis and prepared the manuscript

F.K. supervised the research, provided methodological guidance and revised the manuscript

B.K. prepared the manuscript, reviewed and edited the work and provided methodological guidance

K.M. prepared the manuscript, reviewed and edited the work and provided methodological guidance

W.S. prepared the manuscript, reviewed and edited the work and provided methodological guidance

M.M. supervised the research work, prepared the manuscript, reviewed and edited the work and provided methodological guidance

G.K. collected field samples, helped in statistical analysis and reviewed the manuscript

E.C. conducted data curation, strengthened statistical analysis and reviewed the manuscript

W.S. supervised the research, provided methodological guidance, reviewed the statistical analysis and manuscript

E.N. supervised the research, provided methodological guidance, reviewed the statistical analysis and manuscript, and validated the work

All authors contributed to the reading, writing, reviewing, and approved the final version of the manuscript

## Ethical approval

Ethical approval was granted by Mzuzu University Research Ethics Committee (Approval No: MZUNIREC/DOR/24/83)

## Submission declaration

The authors confirm that this manuscript is original, has not been published previously, and is not under consideration for publication elsewhere. All authors approved the final manuscript for submission.

## Supplemental Tables

**Figure S1.**
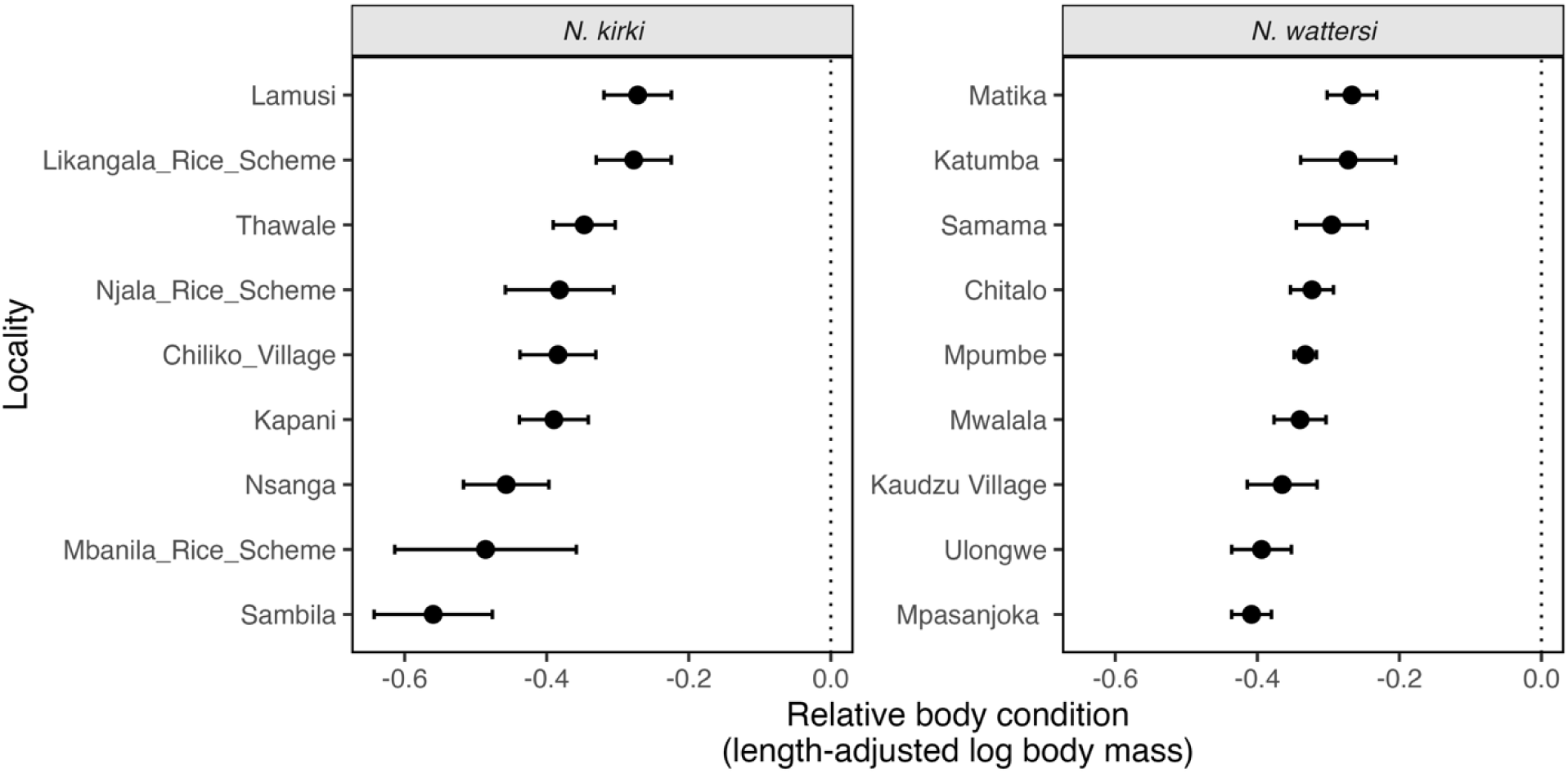
Population variation in relative body condition among localities of *N. kirki* and *N. wattersi*. Points represent estimated marginal means of length-adjusted log body mass derived from analysis of covariance models controlling for body length and sex. Horizontal bars indicate 95% confidence intervals. Localities are ordered by relative body condition within each species. Values closer to zero indicate individuals that were heavier for a given body length, whereas more negative values indicate individuals that were lighter than expected after accounting for body size.

**Figure S2.**
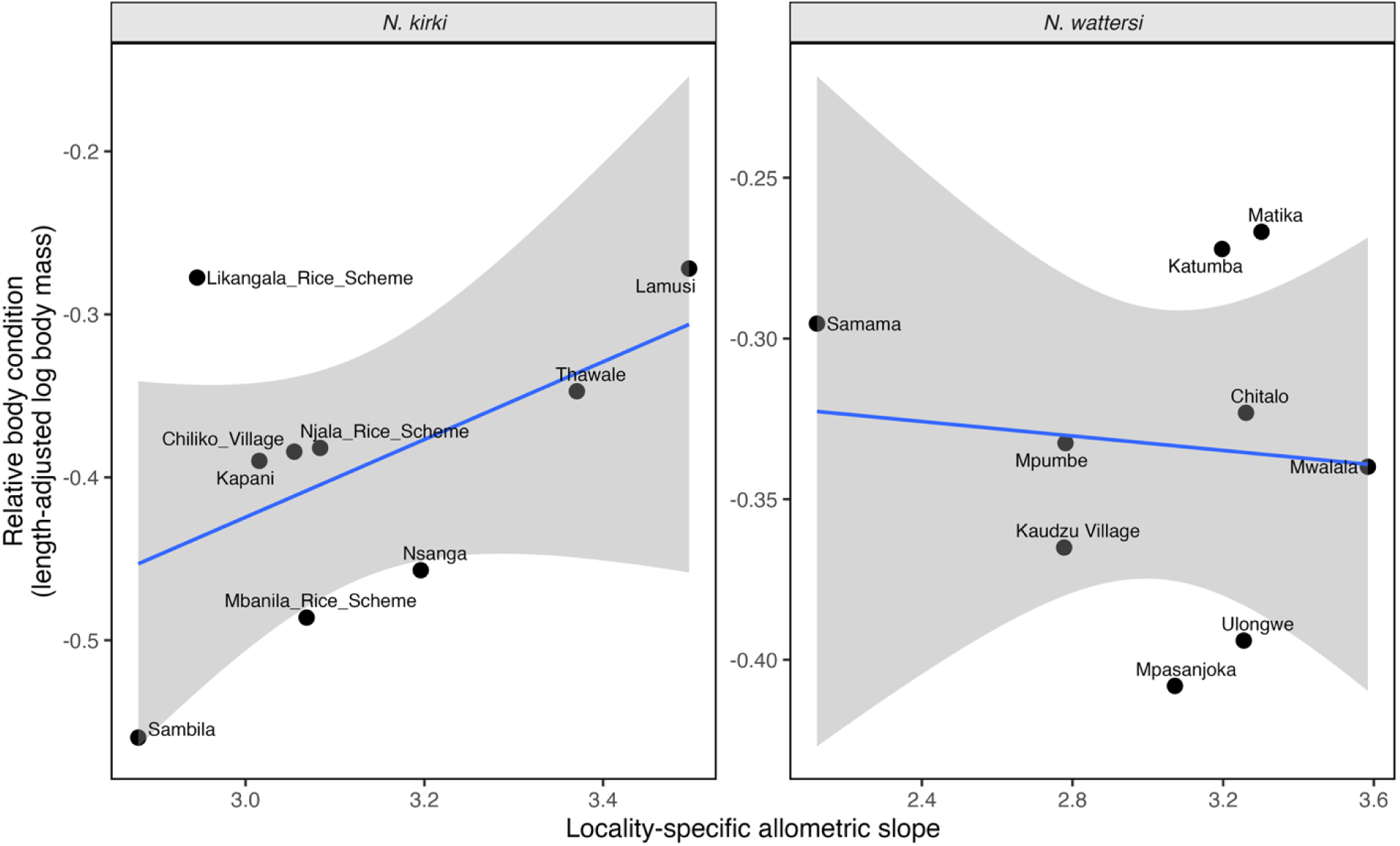
Relationships between population allometric coefficients and relative body condition in *N. kirki* and *N. wattersi*. Points represent localities, with relative body condition estimated as length-adjusted log body mass derived from models controlling for body length and sex. Blue lines indicate fitted linear regressions and shaded regions represent 95% confidence intervals. These analyses were used to evaluate whether populations exhibiting different growth trajectories also differed in average body condition.

**Figure S3.**
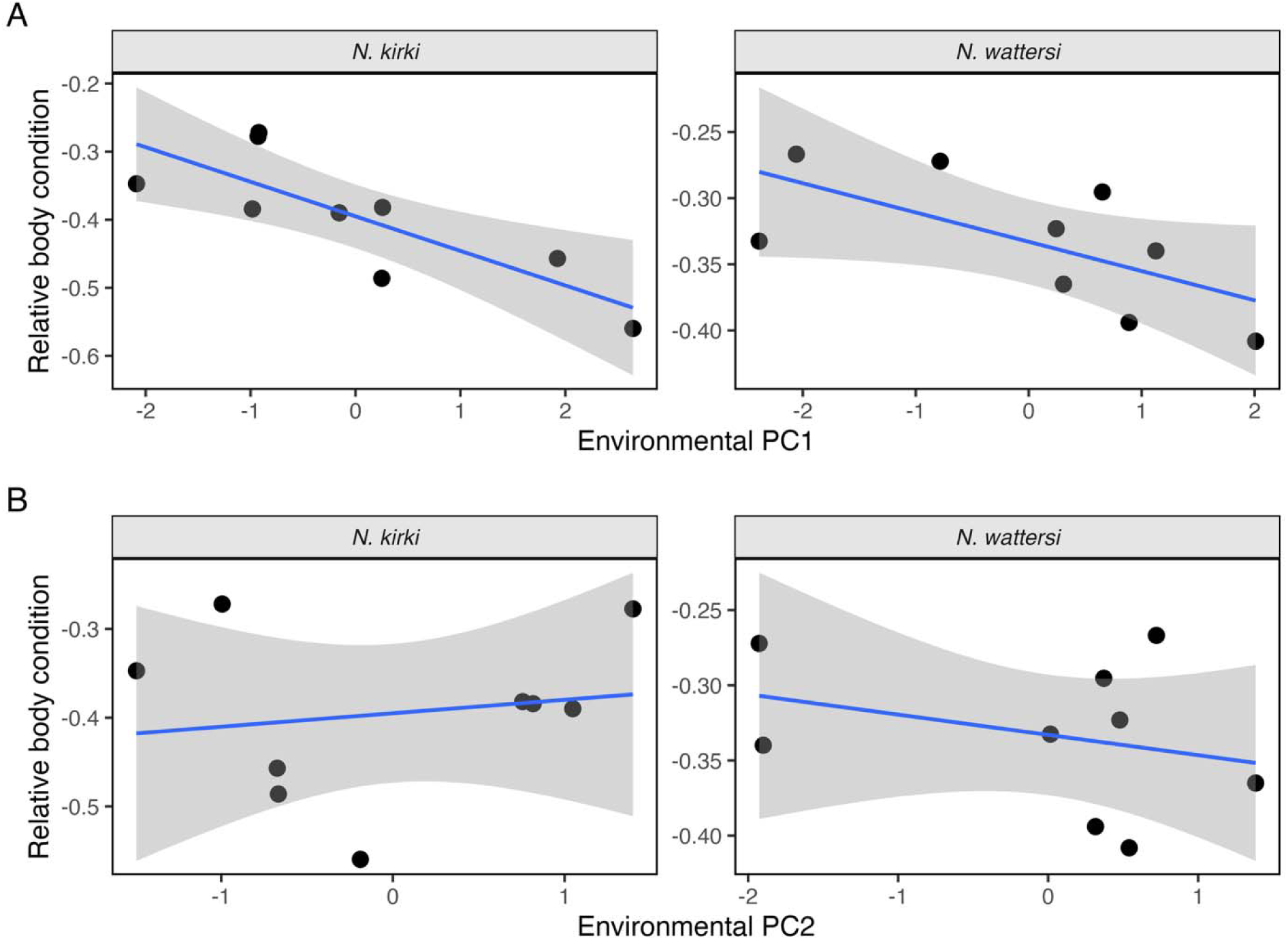
Associations between environmental variation and relative body condition in *N. kirki* and *N. wattersi*. Relative body condition (length-adjusted log body mass) was regressed against the first two environmental principal component axes derived from standardized measurements of temperature, salinity, conductivity, dissolved oxygen, and pH. **(a)** Relationships with PC1 and **(b)** relationships with PC2. Points represent localities, regression lines indicate fitted linear models, and shaded regions represent 95% confidence intervals.

## Supplemental figures

**Table S1.** Relative body condition among populations of *N. wattersi* and *N. kirki*. Estimated marginal means (EMMs) of log-transformed body mass adjusted for body length and sex. Higher values indicate greater relative body condition at a common body length. Confidence intervals are based on the fitted ANCOVA models.

| Species | Population | EMM | Lower 95% CI | Upper 95% CI |
| --- | --- | --- | --- | --- |
| *N. wattersi* | Matika | -0.277 | -0.310 | -0.244 |
|  | Katumba | -0.290 | -0.337 | -0.243 |
|  | Samama | -0.304 | -0.353 | -0.255 |
|  | Mpumbe | -0.326 | -0.341 | -0.312 |
|  | Chitalo | -0.335 | -0.353 | -0.316 |
|  | Kaudzu Village | -0.350 | -0.376 | -0.323 |
|  | Mwalala | -0.352 | -0.378 | -0.327 |
|  | Ulongwe | -0.399 | -0.441 | -0.358 |
|  | Mpasanjoka | -0.403 | -0.419 | -0.387 |
| *N. kirki* | Lamusi | -0.272 | -0.319 | -0.224 |
|  | Likangala Rice Scheme | -0.277 | -0.330 | -0.225 |
|  | Thawale | -0.347 | -0.391 | -0.304 |
|  | Njala Rice Scheme | -0.382 | -0.458 | -0.306 |
|  | Chiliko Village | -0.384 | -0.438 | -0.331 |
|  | Kapani | -0.390 | -0.438 | -0.341 |
|  | Nsanga | -0.457 | -0.517 | -0.397 |
|  | Mbanila Rice Scheme | -0.486 | -0.614 | -0.358 |
|  | Sambila | -0.560 | -0.643 | -0.477 |

